# CaMKII binding to GluN2B flips a β-adrenergic switch from synaptic depression to potentiation

**DOI:** 10.1101/2022.06.23.497296

**Authors:** Olivia R. Buonarati, Matthew E. Larsen, Hai Qian, Johannes W. Hell, K. Ulrich Bayer

## Abstract

Learning, memory and cognition are thought to require forms of synaptic plasticity such as hippocampal long-term potentiation and depression (LTP and LTD), and such plasticity can be modulated by β-adrenergic stimulation with isoproterenol or norepinephrine. For instance, LTP versus LTD is induced by high-versus low-frequency stimulation (HFS versus LFS) but, stimulating β-adrenergic receptors (βARs) enables LTP induction also by LFS. In contrast to HFS-LTP, such βAR-LTP requires signaling by L-type voltage-gated Ca^2+^-channels, not NMDA-type glutamate receptors (NMDARs). Surprisingly, we found that βAR-LTP still required a non-ionotropic NMDAR function: the stimulus-induced binding of the Ca^2+^/calmodulin-dependent protein kinase II (CaMKII) that mediates CaMKII movement to excitatory synapses. In hippocampal neurons, β-adrenergic stimulation with isoproterenol transformed LTD-type CaMKII movement to LTP-type movement, resulting in CaMKII movement to excitatory instead of inhibitory synapses. Additionally, isoproterenol enabled induction of a major cell-biological feature of LTP in response to LTD stimuli: increased SEP-GluA1 surface expression. Like for the βAR-LTP in hippocampal slices, the effects of isoproterenol on CaMKII movement and SEP-GluA1 surface expression involved L-type Ca^2+^-channels. Taken together, these results indicate that isoproterenol transforms LTD stimuli to LTP signals by switching CaMKII movement and GluN2B binding to LTP mode.

**One Sentence Summary:** Buonarati et al. show that β-adrenergic stimulation enables LTP induction in response to LTD stimuli by switching synaptic CaMKII movement to LTP mode.

## INTRODUCTION

Norepinephrine is famous for its participation in the fight or flight response via stimulating β-adrenergic receptors (βARs), but it can also affect memory. Learning and memory are thought to require changes in the number of synaptic AMPA-type glutamate receptors (AMPARs), which can cause long-term changes in synaptic strength such as hippocampal LTP and LTD (Buonarati et al., 2019; Greger and Esteban, 2007; Groc and Choquet, 2020; Herring and Nicoll, 2016; Shepherd and Huganir, 2007). Induction of LTP and some forms of LTD typically require Ca^2+^-influx through NMDA-type glutamate receptors (NMDARs), but with distinct stimulation patterns: Hippocampal LTP is typically induced by brief high-frequency stimulation (HFS; such as 1-4x 1 s at 100 Hz) that causes brief but strong Ca^2+^-stimuli, whereas LTD is typically induced by low-frequency stimulation (LFS; such as 15 min at 1 Hz) that causes weak but prolonged Ca^2+^-stimuli (Bliss and Lomo, 1973; Dudek and Bear, 1992; Yang et al., 1999). Interestingly, β-adrenergic stimulation with isoproterenol enables LTP induction even after low-frequency stimuli (1 to 5 Hz for 15 min) that otherwise instead induce LTD (Huang and Kandel, 2005). Most commonly, such βAR-LTP has been induced with a prolonged theta-frequency tetanus of 5 Hz for 3 min (O’Dell et al., 2015; Thomas et al., 1996), and has also been named for the type of electrical stimulation, i.e. PTT-LTP, as it mimics naturally occurring brain wave patterns (Patriarchi et al., 2016; Qian et al., 2017). In contrast to HFS-induced LTP, this βAR/PTT-LTP is blocked by L-type Ca^2+^ channel inhibition or by conditional Cav1.2 knockout, but is only partially reduced by the glutamate-competitive NMDAR inhibitor APV and is not significantly impaired by the NMDAR pore-blocker MK801 that prevents Ca^2+^-influx through NMDARs (Qian et al., 2017).

We show here that βAR-LTP still requires not only Ca^2+^/calmodulin-dependent protein kinase II (CaMKII) activity but also CaMKII binding to the NMDAR subunit GluN2B. Thus, even though βAR-LTP is not dependent on ionotropic NMDAR signaling (i.e. it does not require Ca^2+^ flux through NMDARs), it is completely dependent on the non-ionotropic scaffolding function of the NMDAR that mediates CaMKII targeting to excitatory synapses. Indeed, we additionally show that β-adrenergic stimulation redirects CaMKII movement from inhibitory to excitatory synapses after LTD stimuli, i.e. the movement normally seen only after LTP stimuli.

## RESULTS

### βAR-LTP requires CaMKII stimulation and it binding GluN2B

In contrast to HFS-induced LTP, the βAR/PTT-LTP that is induced in hippocampal slices by 3 min of 5 Hz stimulation in the presence of 1 μM isoproterenol requires Ca^2+^ influx through the L-type Ca^2+^ channel Cav1.2 but not through the NMDAR (Qian et al., 2017). However, like HFS-LTP, this βAR-LTP was dependent on CaMKII, as it was completely blocked by the Ca^2+^/CaM-competitive CaMKII inhibitor KN93 (Figure 1A,B). Thus, we decided to additionally test if the βAR-LTP also still requires the CaMKII binding to the NMDAR subunit GluN2B that is necessary for normal HFS-induced LTP (Barria and Malinow, 2005; Halt et al., 2012). For this purpose, we used the GluN2B^ΔCaMKII^ mutant mice (Halt et al., 2012) that carry two point mutations in GluN2B (L1298A/R1300Q), which together completely disrupt CaMKII binding (Buonarati et al., 2020; Strack et al., 2000). Notably, the related CaM kinase DAPK1 binds to a partially overlapping site on GluN2B (Goodell et al., 2017; Tu et al., 2010), but DAPK1 binding is completely unaffected by the GluN2B^ΔCaMKII^ mutation (Buonarati et al., 2020). Remarkably, the GluN2B^ΔCaMKII^ mutation completely blocked the βAR-LTP (Figure 1C,D). Accordingly, even though βAR-LTP does not require the ionotropic NMDAR functions that are necessary for HFS-induced LTP, the βAR-LTP is completely dependent on a non-ionotropic NMDAR function: providing a scaffold for binding of CaMKII.

**Figure 1.**
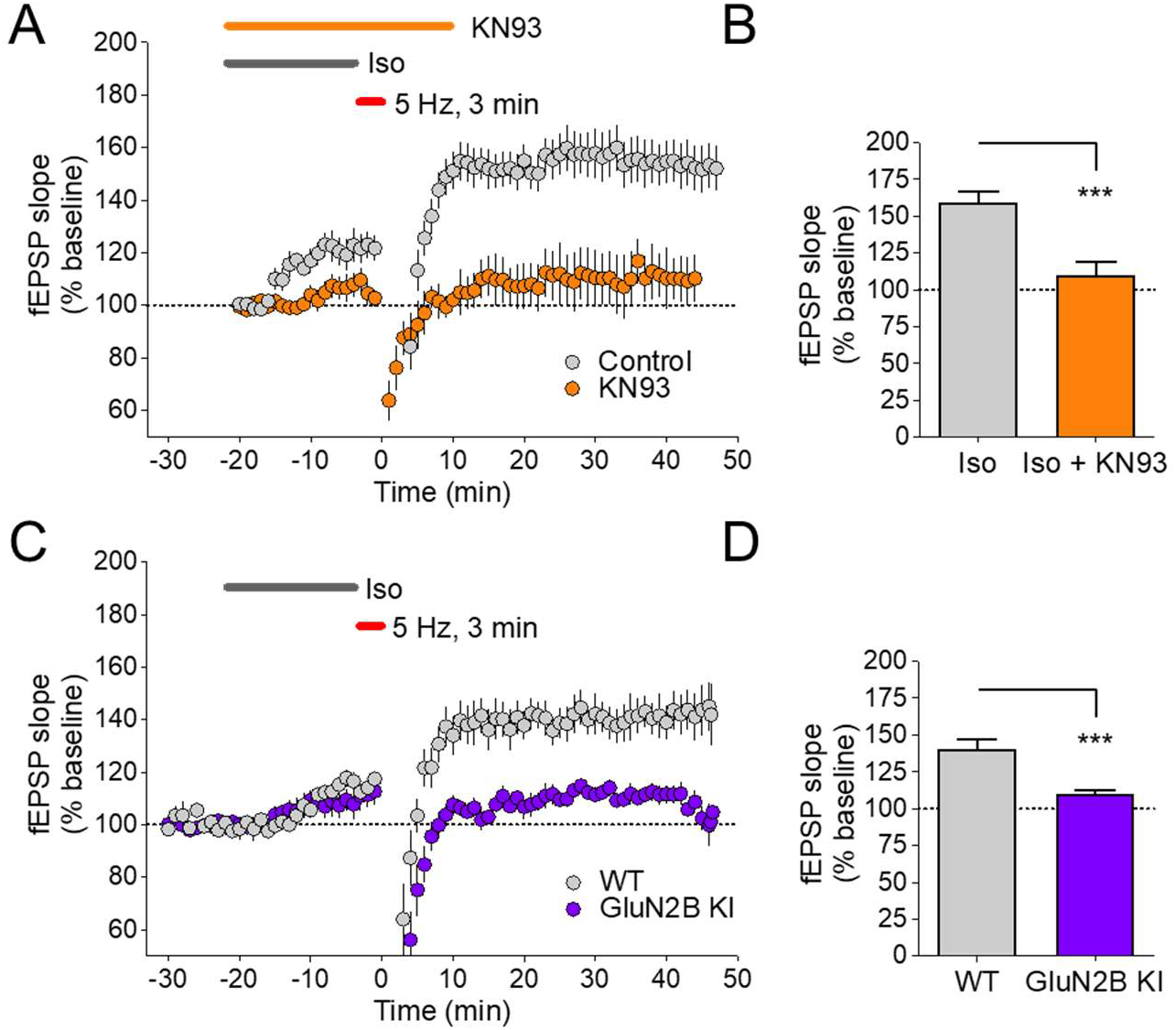
CaMKII binding to GluN2B is required for βAR-LTP. Graphs show fEPSP initial slopes recorded from hippocampal CA1 before and after 3 min, 5 Hz stimulation. Gray bars indicate perfusion with 1 μM isoproterenol (Iso). Quantifications show mean ± SEM. ***p<0.001. (**A**) LTP induced by Iso/5 Hz 3 min is blocked by 10 μM KN-93. (**B**) Potentiation was 159.5 ± 7.4% for Iso and 110.12 ± 9.3% for Iso + KN93 (p<0.001 baseline vs. Iso, 8 mice, 12 slices; ns baseline vs. Iso + KN93, 3 mice, 4 slices). (**C**) LTP induced by iso/5 Hz 3 min stimulation requires CaMKII binding to GluN2B, as potentiation is blocked in slices from GluN2B^ΔCaMKII^ KI mice. (**D**) Potentiation was 140.8 ± 6.2% for slices from wild-type mice and 110.2 ± 2.5% for slices from GluN2B^ΔCaMKII^ KI mice (p<0.001 baseline vs. wild-type, ns baseline vs. GluN2B^ΔCaMKII^ KI; 3 mice, 5 slices per genotype).

### β-adrenergic signaling primes CaMKII for LTP-like synaptic movement upon cLTD stimulation

The Ca^2+^/CaM-induced CaMKII binding to GluN2B mediates the further accumulation of CaMKII at excitatory synapses within dendritic spines in response to LTP-inducing stimuli (Bayer et al., 2001; Halt et al., 2012). By contrast, LTD-inducing stimuli cause CaMKII movement to inhibitory synapses instead (Cook et al., 2019; Marsden et al., 2010). As β-adrenergic stimulation with isoproterenol caused LTP induction in response to stimuli that are otherwise more LTD-related, we decided to determine how isoproterenol may modulate the effects of LTD stimuli on synaptic CaMKII movement. For this purpose, we used intrabody-based simultaneous live-imaging of endogenous CaMKII and two marker proteins for excitatory versus inhibitory synapses, PSD95 and gephyrin (Buonarati et al., 2020; Cook et al., 2019). As expected, CaMKII accumulated at excitatory synapses of hippocampal neurons in response to chemical LTP stimuli (cLTP; 1 min 100 μM glutamate/10 μM glycine) (Figure 2A), but not in response to chemical LTD stimuli (cLTD; 3 min 30 μM NMDA, 10 μM glycine, 10 μM CNQX in the presence of Mg^2+^) (Figure 2B). However, pre-treatment with isoproterenol (1 μM) enabled CaMKII movement to excitatory synapses also in response to cLTD stimuli (Figure 2C). Similar to the βAR-LTP in hippocampal slices (Qian et al., 2017), this isoproterenol-enabled redirection of CaMKII movement was suppressed by the L-type Ca^2+^ channel inhibitor isradipine (10 μM) (Figure 2D,E): Even though some residual movement to excitatory synapses was still observed (Figure 2D), the CaMKII movement was significantly reduced by the addition of isradipine (Figure 2E).

**Figure 2.**
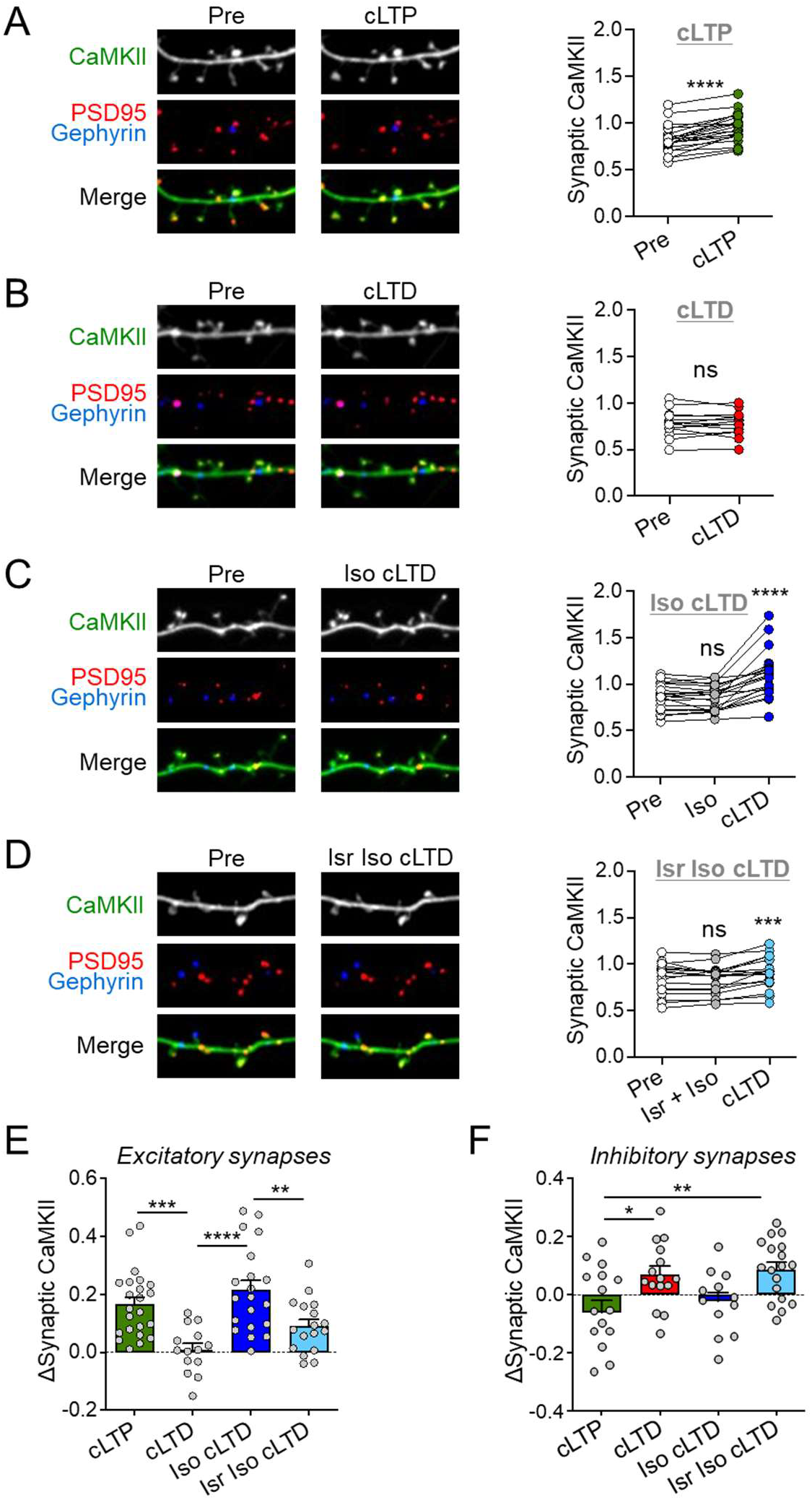
β-adrenergic signaling primes CaMKII for LTP-like synaptic movement after cLTD stimulation. Representative confocal images show endogenous CaMKIIα at excitatory synapses (marked by endogenous PSD-95 in red) and inhibitory synapses (marked by gephyrin in blue) in rat hippocampal neurons (DIV16-18). Quantifications show mean ± SEM. *p<0.05, **p<0.01, ***p<0.001, ****p<0.0001, ns indicates no significance. (**A**) CaMKII moves to excitatory synapses after cLTP (100 μM glutamate, 10 μM glycine, 1 min) (paired two-tailed t-test: pre vs. cLTP ****p<0.0001, n=22 cells). (**B**) No CaMKII movement to excitatory synapses is observed after cLTD (30 μM NMDA, 10 μM glycine, 10 μM CNQX, 3 min) (paired two-tailed t-test: pre vs. cLTD p=0.8348, n=14 cells).(**C**) CaMKII moves to excitatory synapses when cLTD follows pretreatment with isoproterenol (1 μM Iso, 5 min) (repeated-measures oneway ANOVA, Tukey test: pre vs. Iso p=0.8175, pre vs. Iso cLTD ****p<0.0001, Iso vs. Iso cLTD ****p<0.0001, n=20 cells). (**D**) CaMKII movement after Iso cLTD is weakened by isradipine (10 μM Isr, 10 min) (repeated-measures one-way ANOVA, Tukey test: pre vs. Isr Iso p=0.7019, pre vs. Isr Iso cLTD **p=0.0036, Isr Iso vs. Isr Iso cLTD ***p=0.0008, n=17 cells). (**E**) Quantified change in CaMKII movement to excitatory synapses (one-way ANOVA, Tukey test: cLTP vs. cLTD ***p=0.0007, cLTD vs. Iso cLTD ****p<0.0001, Iso cLTD vs. Isr Iso cLTD **p=0.0092, cLTP vs. Isr Iso cLTD p=0.1751, cLTP vs. Iso cLTD p=0.5321, cLTD vs. Isr Iso cLTD p=0.2060). (**F**) Quantified change in CaMKII movement to inhibitory synapses (one-way ANOVA, Tukey test: cLTP vs. cLTD *p=0.0308, cLTP vs. Isr Iso cLTD **p=0.0068, cLTP vs. Iso cLTD p=0.8489, cLTD vs. Iso cLTD p=0.2675, cLTD vs. Isr Iso cLTD p=0.9761, Iso cLTD vs. Isr Iso cLTD p=0.1112).

### β-adrenergic stimulation blocks CaMKII movement to inhibitory synapses upon cLTD stimulation

The reciprocal effect was observed for CaMKII movement to inhibitory synapses, which was significantly greater in response to cLTD-versus cLTP-inducing stimuli (Figure 2F), as expected based on previous studies (Cook et al., 2021; Cook et al., 2019). Adding β-adrenergic stimulation with 1 μM isoproterenol before the cLTD stimulus blocked any movement to inhibitory synapses, but additional inhibition of L-type Ca^2+^ channels with isradipine restored the movement to the same degree as seen after cLTD without β-adrenergic stimulation (Figure 2F; Figure S1). These data indicate that β-adrenergic stimulation can transform LTD-type stimuli to LTP signals by redirecting CaMKII movement to excitatory synapses. Similar to βAR-LTP in hippocampal slices, the β-adrenergic effect on LTP-like CaMKII movement required L-type Ca^2+^ channels.

### Isoproterenol-induced CaMKII-redirection to excitatory synapses requires GluN2B binding

The activity-induced accumulation of CaMKII at excitatory synapses in response to LTP-stimuli requires Ca^2+^/CaM-induced binding to GluN2B (Bayer et al., 2001; Halt et al., 2012). Thus, we decided to test if this is also the case for the redirection of CaMKII movement to dendritic spines that was seen here after β-adrenergic stimulation with isoproterenol. For this purpose, we compared CaMKII movement in neurons from either wild type mice or GluN2B^ΔCaMKII^ mice that have a GluN2B mutation that is CaMKII binding-incompetent (Buonarati et al., 2020; Strack et al., 2000). As expected, CaMKII moved to excitatory synapses in response to cLTP stimuli in hippocampal neurons from mouse, and this movement was blocked by the GluN2B^ΔCaMKII^ mutation (Figure 3A). As also expected, cLTD stimuli did not cause any CaMKII movement to excitatory synapses, neither in wild type nor in GluN2B^ΔCaMKII^ neurons (Figure 3B). However, β-adrenergic stimulation with isoproterenol enabled CaMKII movement in response cLTD stimuli and, as with cLTP stimuli, this movement was abolished by the GluN2B^ΔCaMKII^ mutation (Figure 3C,D). Thus, CaMKII binding to GluN2B is required for both i) the isoproterenol-induced redirection of synaptic CaMKII movement and ii) the βAR-LTP that is otherwise independent of NMDAR current.

**Figure 3.**
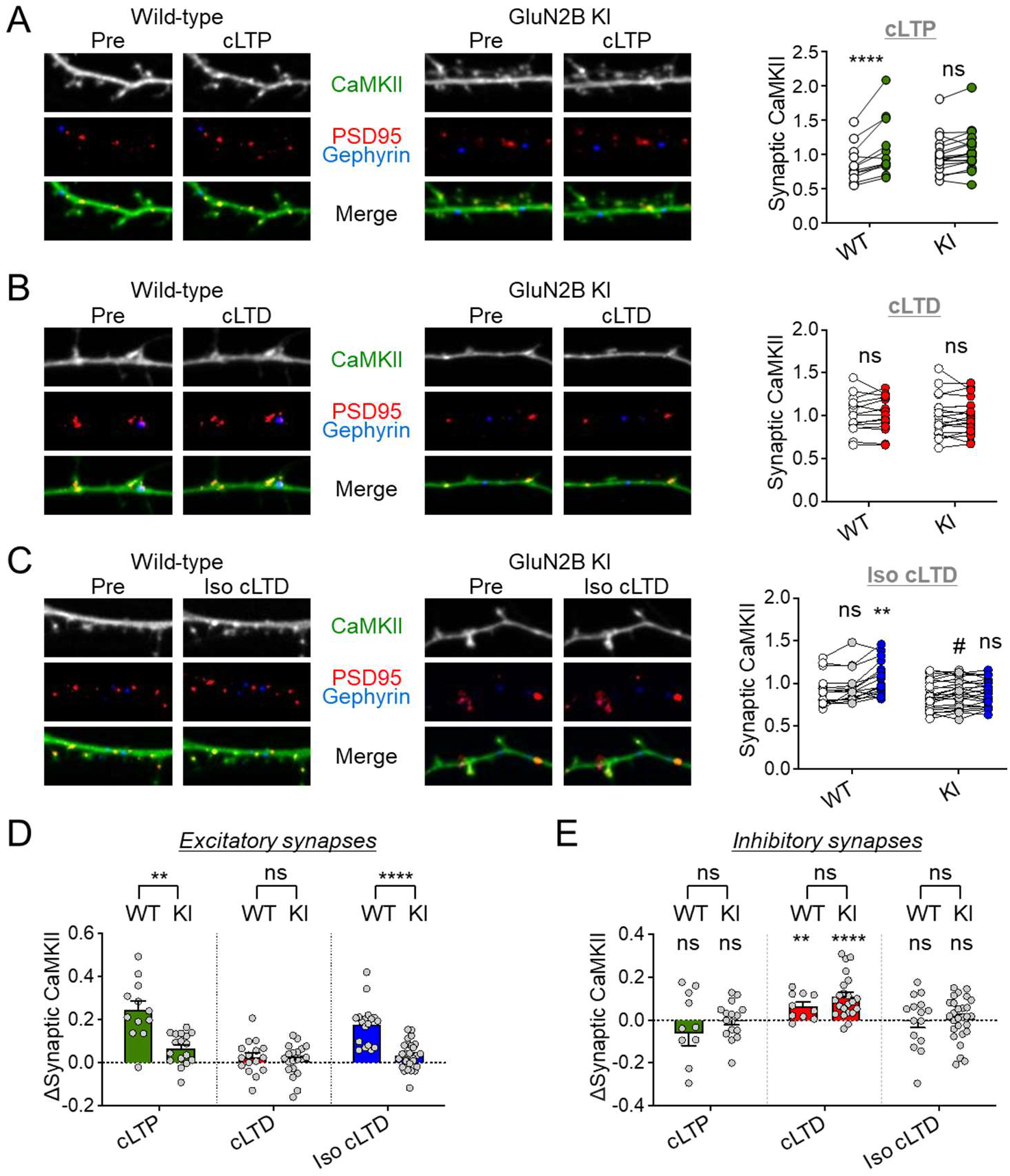
Glu2N2B binding is required for the β-adrenergic switch in synaptic CaMKII movement. Representative confocal images show endogenous CaMKIIα at excitatory synapses (marked by endogenous PSD-95 in red) and inhibitory synapses (marked by gephyrin in blue) in hippocampal neurons (DIV16-18) cultured from wild-type vs. GluN2B^ΔCaMKII^ mice. Quantifications show mean ± SEM. *p<0.05, **p<0.01, ***p<0.001, ****p<0.0001, ns indicates no significance. (**A**) CaMKII moves to excitatory synapses after cLTP (100 μM glutamate, 10 μM glycine, 1 min) in WT but not GluN2B^ΔCaMKII^ KI neurons (two-way ANOVA, Bonferroni’s test: WT pre vs. cLTP ****p<0.0001, n=12 cells; GluN2B KI pre vs. cLTP p=0.0528, n=19). (**B**) No CaMKII movement to excitatory synapses is observed after cLTD (30 μM NMDA, 10 μM glycine, 10 μM CNQX, 3 min) in either WT or KI (two-way ANOVA, Bonferroni’s test: WT pre vs. cLTD p>0.9999, n=15 cells; GluN2B KI pre vs. cLTD p>0.9999, n=20 cells). (**C**) CaMKII moves to excitatory synapses when cLTD is pre-treated with isoproterenol (1 μM Iso, 5 min) only in WT (two-way ANOVA, Bonferroni’s test: pre vs. Iso p=0.1301, pre vs. Iso cLTD ****p<0.0001, Iso vs. Iso cLTD **p=0.0028, n=18 cells). Iso cLTD-induced movement to excitatory synapses is blocked in KI (two-way ANOVA, Bonferroni’s test: pre vs. Iso #p=0.0427, pre vs. Iso cLTD p=0.0839, Iso vs. Iso cLTD p=0.9968, n=25 cells). (**D**) Quantified change in CaMKII movement to excitatory synapses. GluN2B^ΔCaMKII^ KI impairs CaMKII synaptic enrichment compared to WT (two-way ANOVA, Bonferroni’s test: WT vs. KI cLTP **p=0.0025, cLTD p>0.9999, Iso cLTD ****p<0.0001). (**E**) Quantified change in CaMKII movement to inhibitory synapses is not significantly different between WT and GluN2B^ΔCaMKII^ KI (two-way ANOVA, Bonferroni’s test: WT vs. KI cLTP p>0.9999, cLTD p=0.4009, Iso cLTD p>0.9999).

### The GluN2B^ΔCaMKII^ mutation did not cause CaMKII redirection to inhibitory synapses

Whereas the GluN2B^ΔCaMKII^ mutation abolished the LTP-related CaMKII movement to excitatory synapses (Figure 3D), it did not affect CaMKII movement to inhibitory synapses at all (Figure 3E; Figure S2): it neither prevented the LTD-related movement to inhibitory synapses, nor induced such movement in response to LTP-related stimuli. This finding is consistent with the fact that, in contrast to isoproterenol or isradipine, the GluN2B^ΔCaMKII^ mutation does not change the nature of the upstream activation of CaMKII, but only abolishes CaMKII localization at excitatory synapses, which should not affect localization to inhibitory synapses.

### β-adrenergic stimulation enabled SEP-GluA1 surface insertion in response to cLTD stimuli

Our results showed that β-adrenergic stimulation redirects CaMKII to an LTP-like movement in response to cLTD stimuli in hippocampal neurons. This movement was mediated by GluN2B-binding at excitatory synapses, which was required for the βAR-LTP in hippocampal slices. However, in contrast to the βAR-LTP in hippocampal slices, our iso+cLTD stimuli in hippocampal neurons strictly depend on NMDAR signaling, as the cLTD stimulus with 30 μM NMDAR acts by directly activating the NMDAR, without direct activation of other glutamate receptors. Thus, we decided to test the effect of the iso+cLTD stimulus also on increasing the surface expression of the AMPAR subunit GluA1, the hallmark feature of LTP induction. GluA1 surface insertion was measured by expressing SEP-GluA1, i.e. a GluA1 fusion protein with super-ecliptic fluorein, which has quenched fluorescence at the low pH found in vesicles and then increases in fluorescence upon surface insertion (Yudowski et al., 2007). As expected, SEP-GluA1 surface insertion increased significantly only after cLTP but not after cLTD stimuli (Figure 4A,B). However, in presence of 1 μM isoproterenol, even cLTD stimuli triggered an increase in SEP-GluA1 surface (Figure 4C). Without other stimuli, isoproterenol had no effect on SEP-GluA1 surface expression in our neuronal cultures (Figure 4D) (but see (Joiner et al., 2010). Like the βAR-LTP in hippocampal slices and the CaMKII movement in hippocampal neurons, this iso+cLTD-induced SEP-GluA1 insertion in hippocampal neurons was sensitive to inhibition of L-type Ca^2+^ channels: While there was still a trend toward SEP-GluA1 surface insertion, in the presence of 10 μM isradipine, this was no longer significant (Figure 4E; Figure S3). By contrast, the SEP-GluA1 insertion induced by cLTP was not sensitive to such inhibition of L-type Ca^2+^ channels (Figure 4E; Figure S3). Thus, β-adrenergic stimulation of hippocampal neurons with isoproterenol switches both CaMKII movement and GluA1 surface insertion from LTD- to LTP-mode.

**Figure 4.**
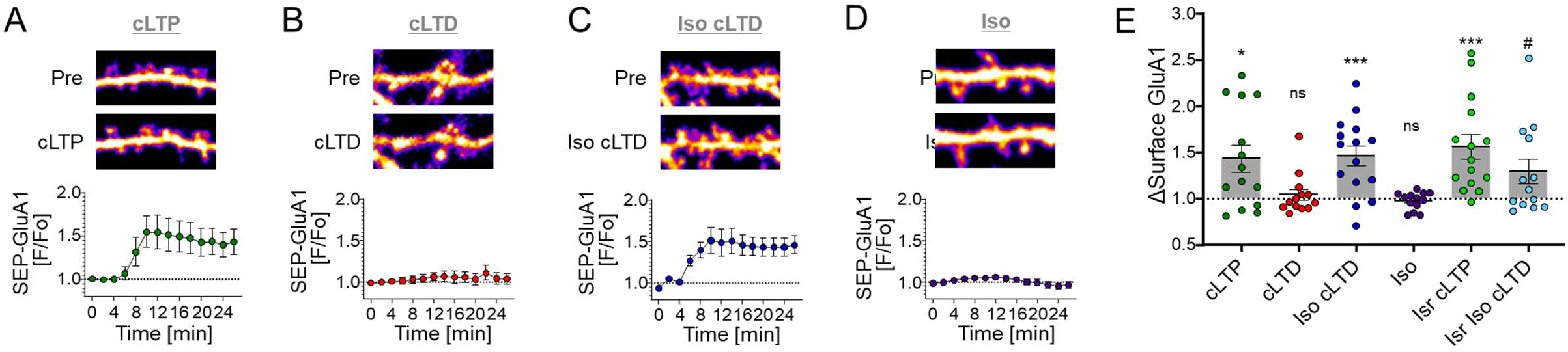
β-adrenergic stimuli allow GluA1 surface insertion in response to LTD stimuli. Representative confocal images show SEP-GluA1 in hippocampal neurons (DIV16-18) cultured from rats. Quantifications show mean ± SEM. *p<0.05, **p<0.01, ***p<0.001, ****p<0.0001, ns indicates no significance. (**A**) cLTP stimuli (100 μM glutamate, 10 μM glycine, 1 min) induced SEP-GluA1 surface insertion. (B) cLTD (30 μM NMDA, 10 μM glycine, 10 μM CNQX, 3 min) stimuli do not induce SEP-GluA1 surface insertion. (**C**) In the presence of β-adrenergic stimulation with isoproterenol (1 μM Iso, 5 min), SEP-GluA1 surface insertion was triggered even by cLTD stimuli. (**D**) On its own, 1 μM isoproterenol had no effect on SEP-GluA1 surface expression. (**E**) Combined bar graphs. Inhibition of L-type Ca2+ channels with isradipine (10 μM Isr, 10 min) did not affect the SEP-GluA1 insertion induced by cLTP, but significantly reduced the insertion in response to iso + cLTD stimuli (one-sample t-test: cLTP *p=0.0111, cLTD ns p=0.545, Iso cLTD ***p=0.0008, Iso ns p=0.3549, Isr cLTP ***p=0.008, Isr Iso cLTD #p=0.056).

## DISCUSSION

Combining electrical 5 Hz stimulation with β-adrenergic stimulation by isoproterenol enables induction of βAR-LTP (also termed PTT-LTP) that depends on L-type Ca^2+^ channels but not on Ca^2+^ influx through NMDARs (Qian et al., 2017). However, this study shows that βAR-LTP still requires a structural, non-ionotropic NMDAR function: synaptic targeting of CaMKII via binding to the NMDAR subunit GluN2B. This binding was absolutely required for βAR-LTP and was responsible for the isoproterenol-induced transformation of activity-induced CaMKII accumulation at excitatory synapses from LTD mode to LTP mode. Notably, βAR-LTP appeared to be even more dependent on CaMKII/GluN2B binding than the HFS-LTP that requires NMDR receptors also for Ca^2+^-influx: Genetic disruption of CaMKII binding to GluN2B reduced HFS-LTP by approximately half (Halt et al., 2012), but abolished βAR-LTP almost completely. This finding also indicates that the most important LTP function of CaMKII binding to GluN2B is not sensitization to local NMDAR Ca^2+^ signals, but rather facilitation of downstream signaling, such as positioning near effector substrates.

The apparent stronger requirement for CaMKII/GluN2B binding in βAR-LTP compared to the “traditional” HFS-LTP could be due to partial compensation in the GluN2B^ΔCaMKII^ mice in which this binding is chronically disabled. Indeed, more acute interventions with CaMKII/GluN2B binding appear to have a stronger effect on such LTP mechanisms (Incontro et al., 2018; Sanhueza et al., 2011). Such compensatory mechanisms for only “traditional” LTP but not βAR-LTP are also consistent with the behavioral findings in the GluN2B^ΔCaMKII^ mice: The modest impairments during the memory consolidation phase in the Morris water maze seen previously (Halt et al., 2012) match the corresponding memory impairment seen when endogenous β-adrenergic signaling with norepinephrine is prevented (Murchison et al., 2004). These modest behavioral effects would be expected from specific impairment of βAR-LTP, whereas general LTP impairment would be expected to lead to a more severe impairment of learning and memory.

HFS-LTP is thought to undergo a developmental switch from PKA- to CaMKII-dependence (Yasuda et al., 2003). By contrast, both kinases appear to be co-required for the βAR-LTP even in adulthood: requirement for phosphorylation of PKA sites on L-type channels in response to β-adrenergic stimulation has been shown previously (Davare et al., 2001; Qian et al., 2017) and our data demonstrate an additional requirement of CaMKII. Both kinases clearly depend on highly localized signaling in order to enable the β-adrenergic conversion of low frequency stimulation to LTP: PKA requires anchoring by AKAP79/150 (Zhang et al., 2013) and CaMKII by GluN2B (shown here). This may provide important mechanisms for how norepinephrine can act as a molecular switch to facilitate acuity during novel, emotionally-charged situations (which would otherwise be less memorable).

Notably, for hippocampal LTD, non-ionotropic functions of the NMDAR have been described to require NMDAR signaling, but not Ca^2+^-flux through the receptor (Gray et al., 2016; Nabavi et al., 2013; Stein et al., 2015). However, this finding remains somewhat controversial (Gray et al., 2016), as others have found block of LTD not only by glutamate-competitive inhibitors, but also by the pore blocker MK801 (Babiec et al., 2014; Coultrap et al., 2014; Dudek and Bear, 1992; Massey et al., 2004; Mulkey and Malenka, 1992; Sanderson et al., 2012). Interestingly, a requirement of CaMKII for LTD has been described not only in studies that found the NMDAR function to be ionotropic (Coultrap et al., 2014) but also in studies that described non-ionotropic NMDAR functions instead (Stein et al., 2021; Stein et al., 2020). However, the CaMKII binding to GluN2B is a non-ionotropic NMDAR function that is specific for LTP: Not only is LTD normal in the GluN2B^ΔCaMKII^ mutant mice (Halt et al., 2012), but LTD even requires specific suppression of the CaMKII binding to GluN2B (Cook et al., 2021; Goodell et al., 2017). Also, the non-ionotropic NMDAR function in LTD still requires NMDAR signaling induced by glutamate binding. Involvement of glutamate binding is less clear for the non-ionotropic scaffolding function in LTP: Even though glutamate-competitive NMDAR inhibitors did not completely block βAR-LTP, they still showed some reduction of βAR-LTP, whereas the channel blocker MK801 had no effect on βAR-LTP in slices at all (Qian et al., 2017). The mechanisms that determine if CaMKII induces LTP or LTD await further elucidation. However, the current study clearly demonstrates that the direction of the LTP-versus LTD-decision by CaMKII is strongly modulated by β-adrenergic signals, both in hippocampal slices and in cultured hippocampal neurons.

In earlier work we found that isoproterenol by itself can stimulate surface insertion of endogenous GluA1 and of ectopically SEP-GluA1 when isoproterenol was applied into the culture medium that contains the B27 or related NS21 supplement (Joiner et al., 2010). In the current work, we replaced the culture medium with ACSF before application of isoproterenol, which clearly prevented isoproterenol by itself to directly induce surface insertion of GluA1. Apparently, the culture medium, which has a complex composition (Chen et al., 2008), contains one or more factors that are not only important for the health of the neurons in culture but also promote GluA1 surface expression so that isoproterenol alone can acutely stimulate it. Defining such factors will be an interesting question for the future, which is now possible to pursue based on our finding of this difference between culture medium and ACSF.

## MATERIALS AND METHODS

### Animal and cell culture models

All animal procedures were approved by the Institutional Animal Care and Use Committee (IACUC) of the University of Colorado Anschutz Medical Campus or of the University of California at Davis; all procedures were carried out in accordance with NIH best practices for animal use. All animals were housed in ventilated cages on a 12 h light/ 12 h dark cycle and were provided ad libitum access to food and water. Mixed sex WT or mutant mouse littermates (on a C57BL/6 background) from heterozygous breeder pairs were used for slice electrophysiology and biochemistry. Mixed sex pups from homozygous mice (P1-2) or Sprague-Dawley rats (P0) were used to prepare dissociated hippocampal cultures for imaging and biochemistry. GluN2B^ΔCaMKII^ knock-in mutant mice were described previously (Goodell et al., 2017; Halt et al., 2012).

### Material and DNA constructs

Material was obtained from Sigma, unless noted otherwise. The expression vectors for the GFP-labeled FingR intrabodies targeting CaMKIIα, PSD-95, and gephyrin were kindly provided by Dr. Donald Arnold (University of Southern California, Los Angeles, CA, USA) as previously characterized (Gross et al., 2013; Mora et al., 2013). As we have described recently (Cook et al., 2019), the fluorophore label was exchanged using Gibson Assembly to contain the following tags in place of GFP: CaMKIIα-FingR-YFP2, PSD-95-FingR-mTurquois, and gephyrin-FingR-mCherry.

### Hippocampal slice preparation

WT and mutant mouse hippocampal slices were prepared using adult mice (8-16 weeks old). Isoflurane anesthetized mice were rapidly decapitated, and the brain was dissected in ice-cold high sucrose solution containing (in mM): 220 sucrose, 12 MgSO_4_, 10 glucose, 0.2 CaCl_2_, 0.5 KCl, 0.65 NaH_2_PO_4_, 13 NaHCO_3_, and 1.8 ascorbate. Transverse hippocampal slices (350 μm) were made using a vibratome (Leica VT 1000A) and transferred into 32°C artificial cerebral spinal fluid (ACSF) containing (in mM): 124 NaCl, 2 KCl, 1.3 NaH_2_ PO_4_, 26 NaHCO_3_, 10 glucose, 2 CaCl_2_, 1 MgSO_4_, and 1.8 ascorbate. All solutions were recovered in 95% O_2_/5% CO_2_ for at least 1 h before experimentation.

### Extracellular field recordings

All recordings and analysis were performed blind to genotype. For electrical slice recording experiments, a glass micro-pipette (typical resistance 0.4 to 0.8 MΩ when filled with ACSF) was used to record field excitatory post-synaptic potentials (fEPSPs) from the hippocampal CA1 dendritic layer in response to stimulation in the Schaffer collaterals at the CA2 to CA1 interface using a tungsten bi-polar electrode. Slices were continually perfused with 30°C ACSF at a rate of 2 mL/min during recordings. Stimuli were delivered every 15 sec and amplified with an Axopatch 2B amplifier, digitized with a Digidata 1320A, and recorded with Clampex 9 (Molecular Devices). Data were analyzed using WinLTP software (Anderson and Collingridge, 2001) with slope calculated as the initial rise from 10 to 60% of response peak. Input/output (I/O) curves were generated by increasing the stimulus intensity at a constant interval until a maximum response or population spike was noted to determine stimulation that elicits 40-70% of maximum slope. Slope of I/O curve was calculated by dividing the slope of response (mV/ms) by the fiber volley amplitude (mV) for the initial linear increase. Paired-pulse recordings (50 ms inter-pulse interval) were acquired from 40% max slope and no differences in pre-synaptic facilitation were seen in mutant slices. Iso LTP was induced by bath application of 1 μM isoproterenol, followed by a train of pulses of a frequency of 5 Hz lasting 3 min. The level of LTP was determined by the average fEPSP initial slope from the 30 min period between 15 and 45 min after the tetanus.

### Hippocampal culture preparation

To prepare primary hippocampal neurons from WT or mutant mice, hippocampi were dissected from mixed sex mouse pups (P1-2), dissociated in papain, and plated at 200-300,000 cells/mL for imaging or 500,000 cells/mL for biochemistry. To prepare rat neurons, hippocampi were dissected from mixed set rat pups (P0), dissociated in papain for 1 h, and plated at 100,000 cells/mL for imaging. At DIV 14-16, neurons were transfected with 1 μg total cDNA per well using Lipofectamine 2000 (Invitrogen), then imaged or treated and fixed 2-3 days later.

### Chemical LTD and LTP stimulation

Chemical LTP (cLTP) was induced with 100 μM glutamate and 10 μM glycine for 1 min. Chemical LTD (cLTD) was induced with 30 μM NMDA, 10 μM glycine, and 10 μM CNQX for 3 min. Iso cLTD was induced by pre-treating cLTD with 1 μM isoproterenol (iso) for 5 min. 10 μM isradipine (isr) was added directly before the iso pre-treatment. Treatments to induce cLTP or cLTD were followed by washout in fresh ACSF.

### Live imaging of CaMKII movement in hippocampal cultured neurons

All images were acquired using an Axio Observer microscope (Carl Zeiss) fitted with a 63x Plan-Apo/1.4 numerical aperture (NA) objective, using 445, 515, 561, and 647 nm laser excitation and a CSU-XI spinning disk confocal scan head (Yokogawa) coupled to an Evolve 512 EM-CCD camera (Photometrics). Experiments were analyzed using Slidebook 6.0 software (Intelligent Imaging Innovations [3i]). During image acquisition, neurons were maintained at 34°C in ACSF solution containing (in mM): 130 NaCl, 5 KCl, 10 HEPES pH 7.4, 20 glucose, 2 CaCl_2_, and 1 MgCl_2_, adjusted to proper osmolarity with sucrose. After baseline imaging and cLTP or cLTD treatment, neurons were imaged 5 min after washout. Tertiary dendrites from pyramidal spiny neurons were selected from maximum intensity projections of confocal Z stacks. To analyze synaptic CaMKIIα, the mean YFP intensity (CaMKIIα) at excitatory (PSD-95) and inhibitory (gephyrin) synapses was quantified. PSD-95 and gephyrin threshold masks were defined using the mean intensity of mTurquois or mCherry plus two standard deviations. Synaptic CaMKIIα was then calculated using the mean YFP intensity at PSD-95 or gephyrin puncta masks divided by the mean intensity of a line drawn in the dendritic shaft. Changes in CaMKIIα synaptic accumulation were determined by dividing the net change in YFP at PSD-95 or gephyrin puncta-to-shaft ratio by the pre-stimulation YFP puncta-to-shaft ratio.

### Live imaging of SEP-GluA1 surface expression in hippocampal cultured neurons

All images were acquired using an Axio Observer microscope (Carl Zeiss) fitted with a 63x Plan-Apo/1.4 numerical aperture (NA) objective, using 488 and 561 nm laser excitation and a CSU-XI spinning disk confocal scan head (Yokogawa) coupled to an Evolve 512 EM-CCD camera (Photometrics). Experiments were analyzed using Slidebook 6.0 software (Intelligent Imaging Innovations [3i]). During image acquisition, neurons were maintained at 34°C in ACSF solution containing (in mM): 130 NaCl, 5 KCl, 10 HEPES pH 7.4, 20 glucose, 2 CaCl_2_, and 1 MgCl_2_, adjusted to proper osmolarity with sucrose. After baseline imaging and cLTP and cLTD treatment, neurons were imaged once every 2 min for 20 min after washout. Tertiary dendrites from pyramidal spiny neurons were selected from maximum intensity projections of confocal Z stacks. To analyze surface GluA1 expression, the mean SEP intensity (GluA1) at distinct SEP puncta was quantified. Changes in GluA1 surface expression were determined by dividing the mean SEP intensity in distinct puncta at 20 min after washout by the pre-stimulation mean SEP intensity.

### Quantification and statistical analysis

All data are shown as mean ± SEM. Statistical significance and sample size (n) are indicated in the figure legends. Data from the imaging experiments were obtained and quantified using SlideBook 6.0 software (3i) and analyzed using Prism (GraphPad) software. To test for parametric conditions, data were evaluated by a Shapiro-Wilk test for normal distribution and a Brown-Forsythe test (3 or more groups) or an F-test (2 groups) to determine equal variance. Comparisons between two groups were analyzed using unpaired, two-tailed Student’s t-tests. Comparisons between pre- and post-treatment images at the same synapse type from the same neurons were analyzed using paired, two-tailed Student’s t-tests. Comparisons between three or more groups were done by one-way ANOVA with Tukey’s post-hoc test. Treatment of the same neuron over time was analyzed as repeated-measures. Comparisons between three or more groups with two independent variables were assessed by two-way ANOVA with Bonferroni post-hoc test to determine whether there is an interaction and/or main effect between the variables.

## Supplementary Materials

Figure S1. CaMKII synaptic enrichment at inhibitory synapses in rat.

Figure S2. CaMKII synaptic enrichment at inhibitory synapses in mouse.

Figure S3. SEP-GluA1 surface insertion in response to cLTP stimuli does not required L-type Ca^2+^-channels

## Acknowledgments

We thank Dr. Steven Coultrap and Ms. Janna Mize-Berge for help with mouse colony maintenance.

## Author contributions

Conceptualization: ORB, JWH, KUB

Investigation: ORB, MEL, HQ Visualization: ORB, MEL, KUB

Funding acquisition: ORB, JWH, KUB

Project administration: KUB

Writing – original draft: ORB, KUB

Writing – review & editing: ORB, MEL, JWH, KUB

## Data and materials availability

All data needed to evaluate the conclusions are present in the paper. Additionally, the datasets generated during this study are available through Mendeley: (put in the link).

## SUPPLEMENTAL FIGURES

**Fig. S1.**
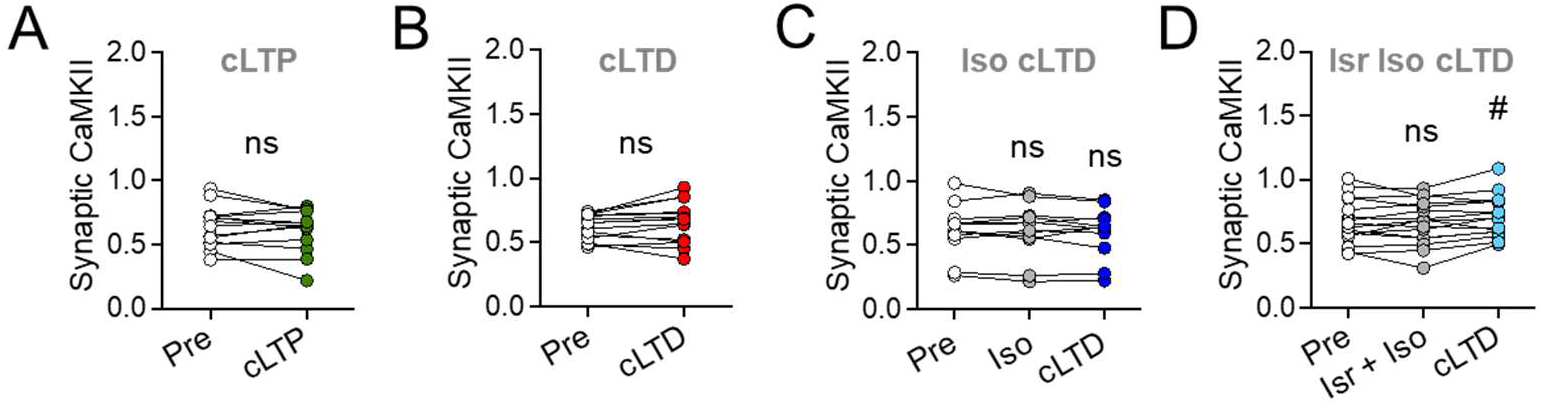
CaMKII synaptic enrichment at inhibitory synapses in rat. Related to Figure 2. Endogenous CaMKIIα at inhibitory synapses (marked by gephyrin) in rat hippocampal neurons (DIV16-18). Quantifications show mean ± SEM. (**A**) No significant CaMKII movement to inhibitory synapses was detected after cLTP (paired two-tailed t-test: pre vs. cLTP p=0.3451, n=14 cells); (**B**) While the CaMKII movement to inhibitory synapses after cLTD was significantly higher than after cLTP (see Fig. 2F), it did not reach significance in a paired test (paired two-tailed t-test: pre vs. cLTD p=0.1462, n=15 cells). By contrast, significance was seen also in paired tests in our previous studies in neurons from both rat and mice (Cook et al., 2021; Cook et al., 2019) and here in mouse neurons (see Supplemental Figure 3B), overall indicating that cLTD stimuli indeed cause CaMKII movement to inhibitory synapses. (**C**) No significant CaMKII movement to inhibitory synapses was detected after Iso cLTD (repeated-measures one-way ANOVA, Tukey test: pre vs. Iso p=0.9717, pre vs. Iso cLTD p=0.6339, Iso vs. Iso cLTD p=0.7709, n=12 cells). In contrast to cLTD withou iso, this movement did not differ from the cLTP condition (see Fig. 2F). (**D**) CaMKII movement to inhibitory synapses after Isr Iso cLTD was not significant in a paired test, but reached a trend (repeated-measures one-way ANOVA, Tukey test: pre vs. Iso Isr p=0.9710, pre vs. Iso Isr cLTD p=0.0739, Isr Iso vs. Isr Iso cLTD #p=0.0605, n=16 cells).

**Fig. S2.**
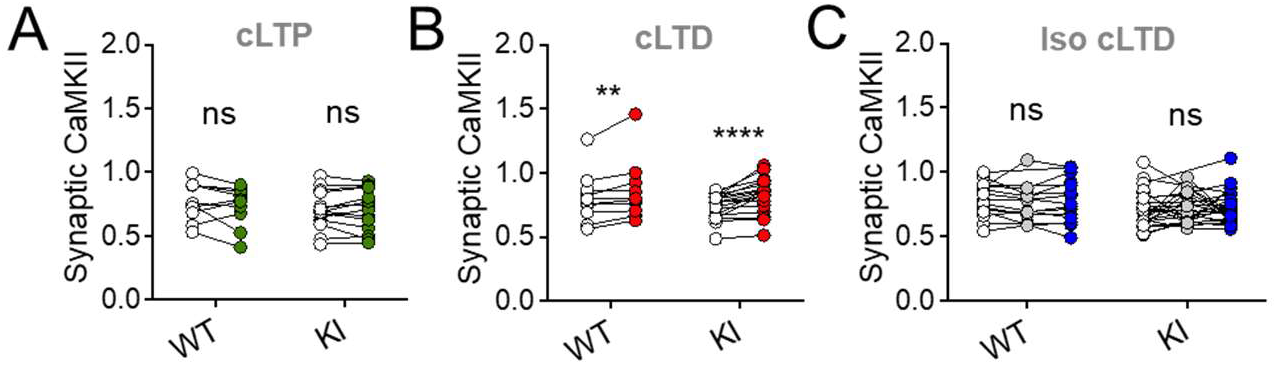
CaMKII synaptic enrichment at inhibitory synapses in mouse. Related to Figure 3. Endogenous CaMKIIα at inhibitory synapses (marked by gephyrin) in hippocampal neurons cultured from WT and GluN2B^ΔCaMKII^ KI mice (DIV16-18). Quantifications show mean ± SEM. (**A**) No significant CaMKII movement to inhibitory synapses is detected after cLTP in either WT or KI (twoway ANOVA, Bonferroni’s test: WT pre vs. cLTP p=0.2552, n=10 cells; GluN2B KI pre vs. cLTP p>0.9999, n=14 cells). (**B**) CaMKII moves to inhibitory synapses after cLTD in both WT and KI (Wilcoxon matched-pairs signed rank test: WT pre vs. cLTD **p=0.0098, n=10 cells; GluN2B KI pre vs. cLTP ****p<0.0001, n=18 cells). (**C**) No significant CaMKII movement to inhibitory synapses is detected after Iso cLTD (repeated-measures one-way ANOVA, Tukey test: WT pre vs. Iso p=0.7123, pre vs. Iso cLTD p=0.9988, Iso vs. Iso cLTD p=0.4232, n=15 cells; KI pre vs. Iso p=0. 9083, pre vs. Iso cLTD p=0.8254, Iso vs. Iso cLTD p=0.9977, n=25 cells).

**Fig. S3.**
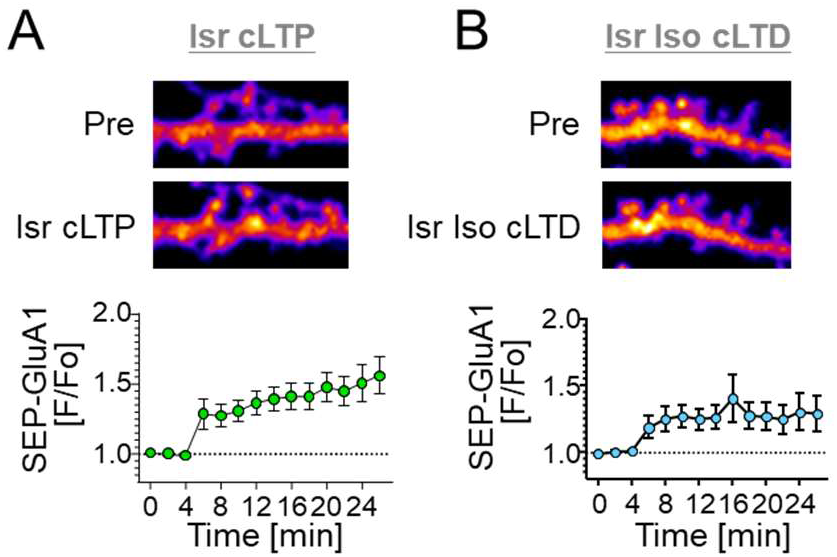
S3. SEP-GluA1 surface insertion in response to cLTP stimuli does not required L-type Ca^2+^-channels, as it is not blocked by 10 μM isradipine. Related to Figure 4. Representative confocal images show SEP-GluA1 in hippocampal neurons (DIV16-18) cultured from rats. Quantifications show mean ± SEM. (**A**) Inhibition of L-type Ca^2+^ channels with isradipine did not affect the SEP-GluA1 insertion induced by cLTP. (**B**) Inhibition of L-type Ca^2+^ channels with isradipine reduced SEP-GluA1 insertion induced by Iso cLTD.

